# An endophytic bacteria (*Pseudomonas aeruginosa*) strain translocated and protected tomato (*Solanum lycopersicum* L.) plants from its parasitic weed *Phelipanche aegyptiaca*

**DOI:** 10.1101/842757

**Authors:** Lilach Iasur Kruh, Jacline Abu-Nassar, Ofir Lidor, Radi Aly

## Abstract

*Phelipanche aegyptiaca* is an obligate holo-parasitic weed lacking a functional photosynthetic system, which subsists on roots of a wide range of host crops, causing severe losses in yield quality and quantity. The parasite and its host are connected through their vascular system, forming a unique ecological system that enables the exchange of various substances. In a previous study, it was suggested that endophytic bacteria, which naturally inhabit the internal tissues of plants, can also be transmitted from the parasitic weed to its host and *vice versa*.

In the current study, we investigate the characteristics of a previously isolated *Pseudomonas* sp. *PhelS10* strain, using both biochemical and molecular methods. Our results revealed that production of *Pseudomonas aeruginosa* quinolone signal (PQS) was 2.1 times higher than that of the standard *Pseudomonas aeruginosa* strain (PAO1), which contributed to a 22% higher biofilm formation capability. *PhelS10* strain was detected in the xylem of tomato plants using FISH analysis. In addition, *PhelS10* strain was found in the parasitic weed’s inner tissues, confirming the hypothesis that endophytic bacteria traffic between the plant host and its parasitic weed.

## Introduction

Broomrapes (*Phelipanche / Orobanche* spp.) are known agricultural pests that attack the roots of many economically important crops throughout the semiarid regions of the world. There are more than 100 species of these obligate holoparasites, including *Phelipanche aegyptiaca, Phelipanche ramosa, Orobanche Cumana, Orobanche minor, Orobanche cernua, and Orobanche crenata* (Parker and Riches, 1993). Parasitic weeds do not possess functional roots, and even though their genome contains photosynthesis genes, they cannot produce organic compounds from CO_2_ (Joel et al., 2006). Instead, they develop special intrusive organs (haustoria) that penetrate crop roots, directly connecting them to the vascular system of the crop plants that serve as their hosts (Westwood, 2000; Joel & Portnoy, 1998). Each mature broomrape produces tens of thousands of seeds, which can remain viable in the soil for many years. The seeds germinate only after receiving a chemical stimulus from a neighboring host root (reviewed in Bouwmeester et al. 2003; Yoder 1999; Koltai and Prandi 2019). Subsequently, the parasite develops an haustorium followed by a tubercle, in which storage reserves accumulate at the expense of host photosynthesis. Eventually, a shoot develops from the tubercle; this shoot emerges above soil, flowers, and carries seeds (Parker and Riches 1993). Broomrapes are the worst root parasite in Israel and in the Mediterranean basin, posing a serious and continual threat to the production of a number of agronomic and horticultural crops, such as tomato, carrot and sunflower. *P. aegyptiaca* in Israel is regarded the most serious pest in vegetables and field crops, with estimated annual damage reaching $10 million. In the Middle East, annual food crop losses from parasitic weeds can be conservatively estimated at $1.3 billion to $2.6 billion (Joel et al., 2013).

A wide variety of parasitic weed control methods have been applied in an attempt to control broomrape (Joel et al., 2006; Aly et al., 2009; Aly, 2007; Cochavi et al., 2016), most of which are based on chemical sprays that can be windborne and are toxic to non-target plants (Aly, 2007). Therefore, there is a need to find alternative solutions to reduce plant-plant parasitization.

The search for agricultural-aiding microorganisms is an emerging strategy to tackle many plant pathogens (Hallmann et al., 1997; Compant et al., 2005; Rosenblueth and Martínez-Romero, 2006; Ryan et al., 2008; Narayanasamy, 2013). It is also a more environment-friendly approach compared to the usage of pesticides in agriculture, as researchers have demonstrated the existence of residual chemicals in agricultural crops, which thereby affect the whole food chain (Ekström et al., 2011; Maheshwari et al., 2013).

Endophytic bacteria that are located in the inner tissue (mostly the parenchyma, intercellular space and vascular tubes) of plants and enhance their host fitness have emerged as effective biocontrol agents (Hardoim et al., 2015; Rosenblueth and Martínez-Romero, 2006; Compant et al., 2010). Recent studies have shown that various bacteria originating from the soil and host plants can suppress parasitic weeds (Joel and Gressel, 2013). For example, the bacterium *Azospirillum brasilense* inhibits seed germination and radical elongation in the broomrape *Phelipanche aegyptiaca*, while *Pseudomonas fluorescens* reduces both the quantity and biomass of the broomrape *Orobanche foetida* (Zermane et al., 2007). The bacterium *Rhizobium* spp. reduces not only seed germination of *O. foetida* but also the number of tubercles on its host’s (chickpea) roots (Hemissi et al., 2013).

In our laboratory, we isolated *Pseudomonas* strain *PhelS10*, obtained from the host plant tomato, which was able to reduce parasitic seed germination *in vitro* and tubercle (*P. aegyptiaca*) development *in planta* (Iasur Kruh et al., 2017). In the current study we further examine this isolate by characterizing its phylogenetic identity and other traits that may facilitate its ability as a biocontrol agent.

## Material & methods

### Bacterial strains and growth conditions

*PhelS10* strain (Prof. Radi Aly lab, ARO) and *Pseudomonas aeruginosa PAO1* wild type strain (obtained from Prof. Steinberg Doron’s laboratory, The Hebrew University of Jerusalem) that serves as a standard strain, were used in order to examine different traits known in the Pseudomonadaceae bacterial family. Bacterial cultures were kept in frozen stocks supplemented with a final 30% glycerol (v/v) at 80°C. These two isolates were grown overnight in LB (Luria Broth, Sigma, Israel) at 37°C and 220 rpm. Starters were grown to turbidity OD=0.1 (600nm); then, the starters were used to inoculate the necessary volume of broth culture at a ratio of 1:100 (V:V) in an Erlenmeyer flask.

### Phylogenetic analysis

In order to characterize the *P. aeruginosa PhelS10* strain phylogenetically, we used three different genes that are known to exist in the Pseudomonadaceae family using the following primers (Curran et al., 2014):

acetyl-coenzyme A synthetase-*acsA* (ACCTGGTGTACGCCTCGCTGAC GACATAGATGCCCTGCCCCTTGAT and GCCACACCTACATCGTCTAT GTGGACAACCTCGGCAACCT)

GMP synthase-*guaA* (CGGCCTCGACGTGTGGATGA GAACGCCTGGCTGGTCTTGTGGTA and AGGTCGGTTCCTCCAAGGTC TCAAGTCGCACCACAACGTC)

phosphoenol pyruvate synthase-*ppsA (*GGTCGCTCGGTCAAGGTAGTGG GGGTTCTCTTCTTCCGGCTCGTAG and GGTGACGACGGCAAGCTGTA TCCTGTGCCGAAGGCGATAC).

One colony of the isolate was picked and resuspended directly into a PCR mixture for amplification of each of the gene fragments (direct colony PCR). The reaction (25 μL) contained: 10 μL APEX 2XRedTaq Mix (Genesee Scientific, USA), 5 pmol of each primer (see above), 12.5 μL DDW and 1 μL DNA as template. The PCR procedure was as follows: DNA was denatured at 95°C for 5 min, followed by 30 cycles at 95°C for 30 sec each, 58°C for 30 sec and 72°C for 1 min, followed by 5 min at 72°C. The PCR product was sequenced from both ends by Hy-labs (Rehovot, Israel) and a consensus sequence was constructed.

### Phylogenetic tree

Sequences of these three genes were aligned to NCBI nt database using Blastn algorithm. Similar sequences from complete genome of other Pseudomonas species were selected for further analysis. For each gene, multiple alignment was preformed using Muscle algorithm in MEGA (version 7.0.2). Alignments were truncated to include and all genes were concatenated to produce a single alignment (3415 bp). Then, maximum likelihood tree was calculated in MEGA.

### (Pseudomonas Quinolone Signal) PQS Quantification and biofilm formation

PQS production was measured by standard protocol (Lidor et al. 2015); briefly, *P. aeruginosa* (*PhelS10)* 20 h cultures were extracted with acidified ethyl acetate. The cultures were calibrated to an absorbance value of 1.0 at A_595_. Extracts were dried, re-suspended in 1:1 ethyl acetate-acetonitrile, and separated by thin-layer chromatography (TLC). The extract was spotted onto the TLC plate as follows: 7 μl of bacterial extract or 5 μl of 20 μM PQS synthetic standard (Sigma). PQS quantity was then determined by exposing resolved TLC plates to 325 nm UV light. Each bacterium was examined in triplicates.

Biofilm formation assay was based on a known crystal violet (CV) staining protocol (Peeters et al., 2008); shortly, a 2 ml of bacterial culture grown overnight at 15 ml tube, 28°C with shaking, was poured and gently rinsed with saline. The remaining bacteria that adhered to the tube were incubated with 2 ml 0.1% (w/v) CV solution for 10 min, followed by 10 min incubation in 2 ml of absolute ethanol. 850 μl from each tube were transferred to a cuvette and measured in a Lab-kits ST-VS-722 model spectrophotometer (U-Therm, HN, Chaina) at OD_595_. Each bacterium was examined in triplicates.

### Plant material

Tomato plants (*Solanum lycopersicum* L. cv T-5) were used as the host plants for the assays. Plants were grown in commercial potting soil (light-medium clay containing 63% sand, 12% silt, and 22% clay, Gal-Marketing Ltd, Israel) and kept at controlled conditions: 25-28°C under cool white neon fluorescent illumination (14/10 h light/dark cycle) at approximately 200 μmol m^−2^ s^−1^ light intensity; the plants were watered as needed. In order to have tomato seedlings with fresh and young roots, seedlings were grown for 18 days, after which the root system was removed, and shoots were dipped in sterile water and planted in soil for ten days until young roots emerged.

*P. aegyptiaca* parasite seeds were collected from an infested tomato field at Qiryat Shemona (Northen Israel) and were kept at 8°C until use. Seeds were surface sterilized (2 min in 70% ethanol and 10 min in 0.6% sodium hyper chloride solution followed by double DDW washes) before the parasitism assay (see below).

### Parasitism assay using *In vitro polyethylene bag system (PE)* (Aly *et al*., 2009)

Ten days after refreshing their roots, tomato seedlings were dipped for 10 min in *P. aeruginosa* (*PhelS10*) culture (grown for 24 h in Lysogeny Broth [LB, Difco]) media adapted to 0.3 of optical density at 600nm (OD_600_) and allowed to dry for 1 h. Then the roots of the tomato plants were surface sterilized to ensure that only endophytic bacteria will be examined. In addition, tomato roots that were dipped in sterile water were used as controls. These roots (either containing *PhelS10* or control) were placed in a polyethylene bag, and *P. aegyptiaca* seeds were dispersed on them. Polyethylene bags were placed in growth chambers under controlled conditions: 25-28°C under cool white neon fluorescent illumination (14/10 h light/dark cycle). Eight days post-application of *P. aegyptiaca* seeds to the host roots, 10 ml of 2 mg L^-1^ of the synthetic plant hormone germination stimulant GR24 were added to each bag to inspire broomrape infection. Two separate experiments were conducted and each treatment included six repeats.

### Fluorescence in situ hybridization (FISH) analysis

The samples of plant tissues (tomato and broomrape that were inoculated with *PhelS10* strain/tomato and broomrape that were not inoculated with *PhelS10* strain/control tomato without the parasitic weed) were taken one-month post-inoculation. Roots of the host plant and tubercle of the parasitic weeds were manually cut longitudinally and subjected to the FISH protocol as previously described (Sakurai et al. 2005). Briefly: the samples were collected directly into Carnoy’s fixative (chloroform: ethanol: glacial acetic acid, 6 : 3 : 1) and fixed overnight, after which the samples were bleached by exposure to 6% of H_2_O_2_ (in ethanol) and incubated for 2 days. After that, the peroxide was replaced with 100% ethanol until the probe hybridization stage. The hybridization of the samples was conducted with hybridization buffer (20 mM Tris-HCl [pH 8.0], 0.9 M NaCl, 0.01% sodium dodecyl sulphate, 30% formamide) containing fluorescent probes (10 pmol ml^−1^). A specific probe was used for *Pseudomonas aeruginosa* strain detection-*pqsA*-cye3 gene 3’CCCGATACCGCCGTTTATCA-5’-cye3 (Pittaya and Khaemaporn, 2014). The probe was hybridized 24 h prior to analysis and finally washed once in DDW, whole mounted on a microscope slide and visualized under an IX81Olympus FluoView500 confocal microscope (Olympus, Tokyo, Japan). Specificity of the detection was confirmed using a noprobe control, and plants that were not inoculated with *PhelS10 strain* served as a negative control. All treatments were conducted in six repeats.

The authors declare no conflict of interest.

## Results and discussion

In a previous study, *PhelS10* strain that was isolated from tomato was identified as *P. aeruginosa*. This bacterium was suggested as a potential biocontrol agent against broomrape, showing reduction of broomrape’s seed germination *in vitro* and broomrape’s tubercle development *in planta* by 70% and 66%, respectively (Iasur Kruh et al., 2017). For this reason, we further examined *PhelS10* strain phylogenetic identity. Fig. 1 presents a phylogenetic tree that combines data from three different gene sequences that represent a multilocus sequence scheme that was developed for *Pseudomonas aeruginosa* typing (Curran et al., 2014): acetyl-coenzyme A synthetase (acsA), phosphoenol pyruvate synthase (ppsA) and GMP synthase (guaA). The *PhelS10* strain was found to be closest to *Pseudomonas aeruginosa* PA96, which is a clinical strain isolated in Guangzhou, China (Déraspe et al., 2014). However, *PhelS10* was also phylogenetically close to strain M18, which was isolated in Shanghai from a sweet melon rhizosphere and proven to have effective biocontrol activity protecting plants against various phytopathogens (Ge et al., 2007). These strains, similar to each other phylogenetically but different in their biochemical abilities, demonstrate the wide diversity in the Pseudomonadaceae family. In the current study, *PhelS10* strain was isolated from an agricultural crop and examined against the pathogenic weed broomrape. However, the close proximity of *PhelS10* to *Pseudomonas aeruginosa* PA96 stresses the need to examine its pathogenicity as part of the risk assessment process, before it can be used as a biocontrol agent.

**Fig. 1.**
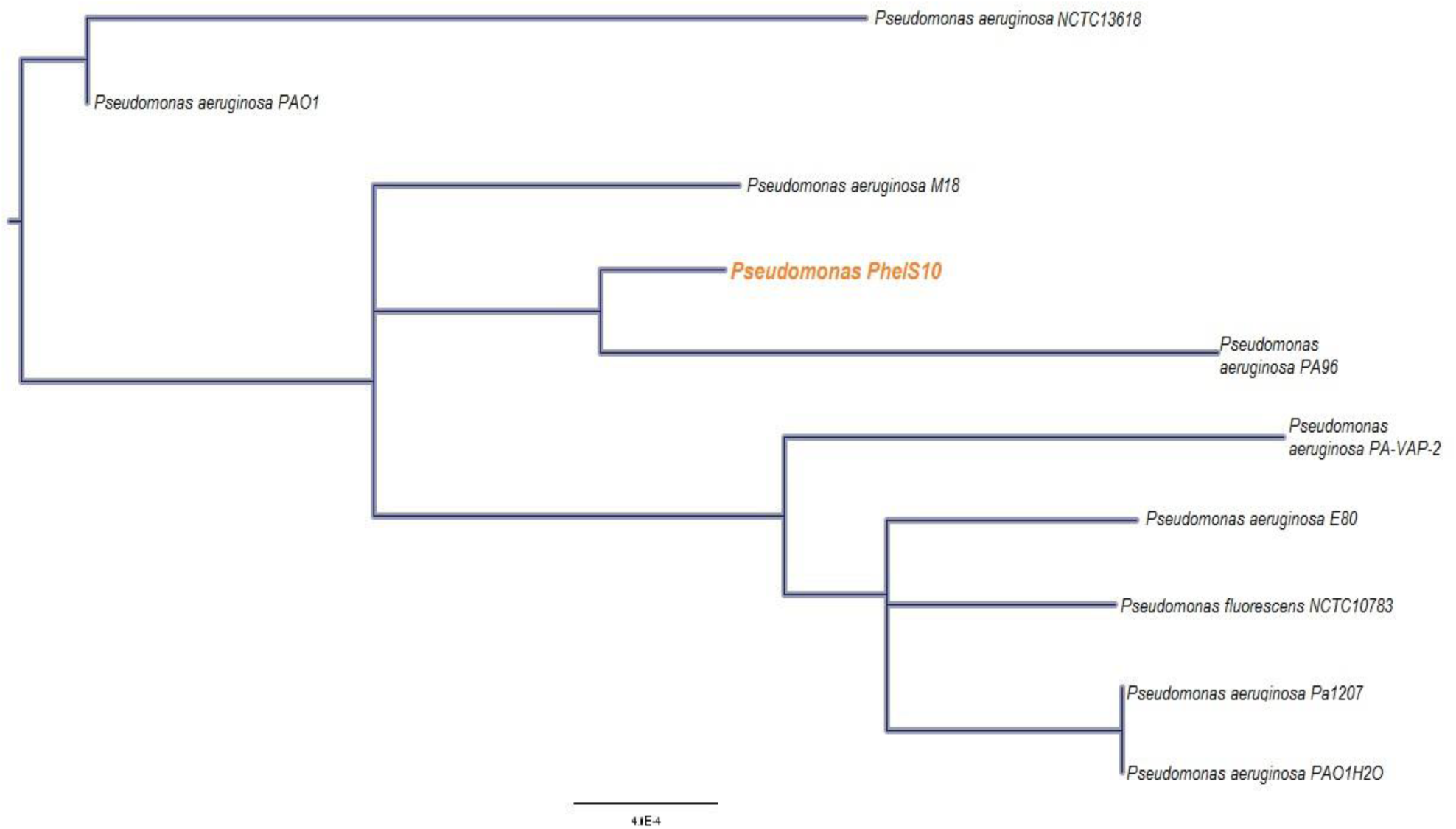
Phylogenetic tree established by multilocus sequence typing scheme for *Pseudomonas aeruginosa*: acetyl-coenzyme A synthetase (acsA), phosphoenol pyruvate synthase (ppsA) and GMP synthase (guaA). A maximum likelihood tree was calculated using gamma distribution rate among sites. Bar indicates rate of substitution per site.

After establishing its phylogenetic identity, *Pseudomonas PhelS10* was examined for its ability to secret a quorum-sensing signal (PQC) as well as to produce biofilm. In comparison to standard *Pseudomonas aeruginosa* strain (PAO1), *PhelS10* produced 2.1 times higher PQS. These signaling systems create a global regulatory network and are believed to regulate the expression of up to 12% of the genome of species belonging to *P. aeruginosa*, including genes related to biofilm production (Lin et al., 2018). Indeed, *PhelS10* produced 22% more biofilm than strain PAO1 (Fig 2). These results indicate that *PhelS10* strain comprises traits that can help it inhabit different environments using biofilm attachment mechanisms. Actually, endophytic bacteria are known to form biofilm as part of their establishment in the intercellular spaces of plant tissues, and this ability was found to affect bacteria-plant relationships (Morris and Monier, 2003).

**Fig. 2.**
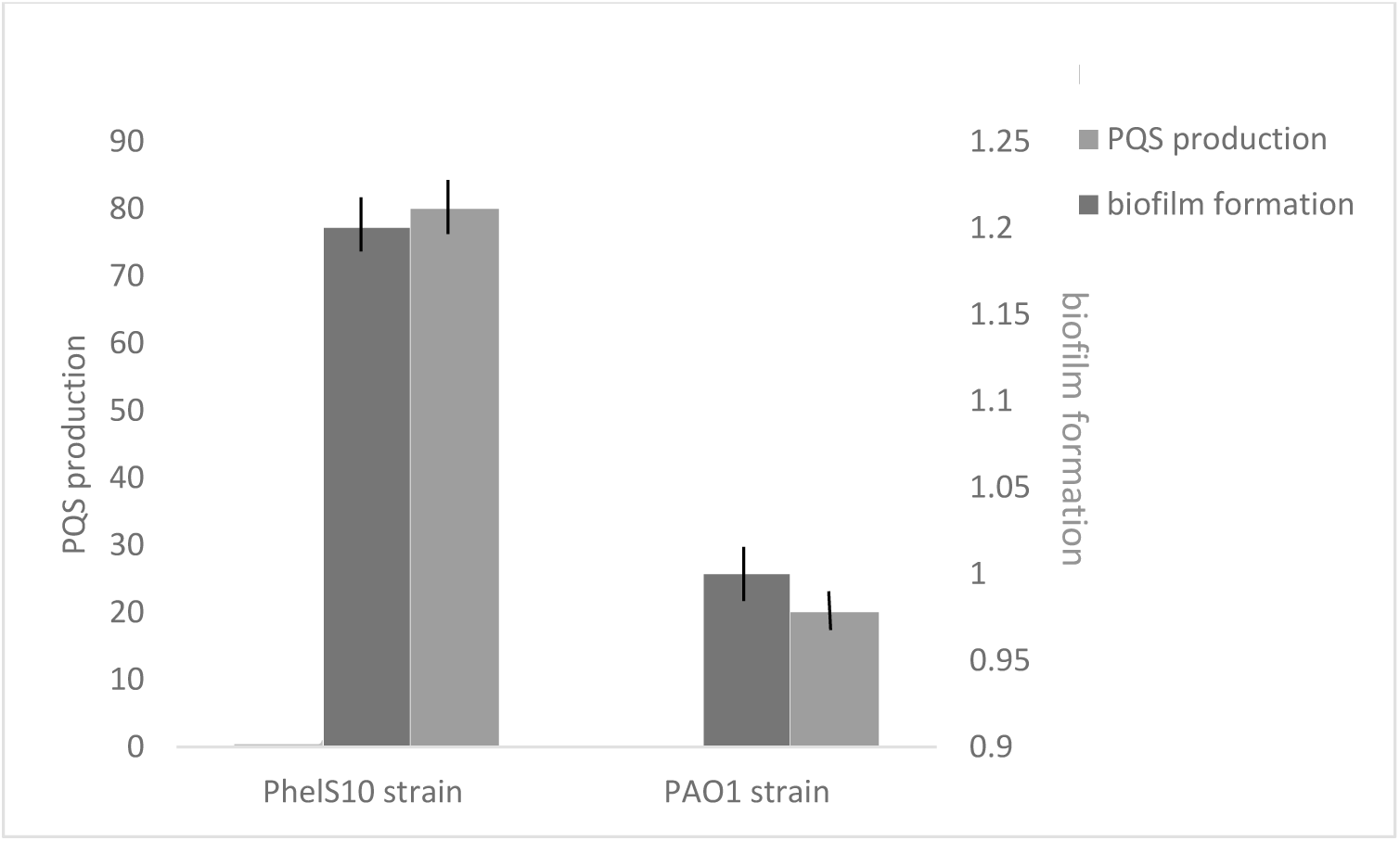
Biofilm formation and PQS production in two *Pseudomonas* strains as measured by standard protocol. The results are an average of three repeats. Bars represent Standard error bars are shown.

Several studies have shown that the passage of endophytic bacteria may be facilitated through the vascular tubes of plants (Hardoim et al., 2015; Rosenblueth and Martínez-Romero, 2006; Compant et al., 2010). In the current study, when tomato roots were augmented with *PhelS10* strain, it penetrated the plant tissue and inhabited the xylem (Fig. 3D). Since the *P. aegyptiaca* tubercle is composed mostly of parenchyma cells which are traversed by both xylem and phloem (Joel et al., 2013), the xylem can serve as a potential transmission route between the parasitic weed and its host, through the haustorium bridge. Indeed, a signal of *PhelS10* strain was detected in the inner tissue of broomrape that was connected to a tomato plant inoculated with this strain (Fig. 3D). No signal was found in the un-inoculated tomato and its broomrape (Fig. 3 A+C).

**Fig. 3.**
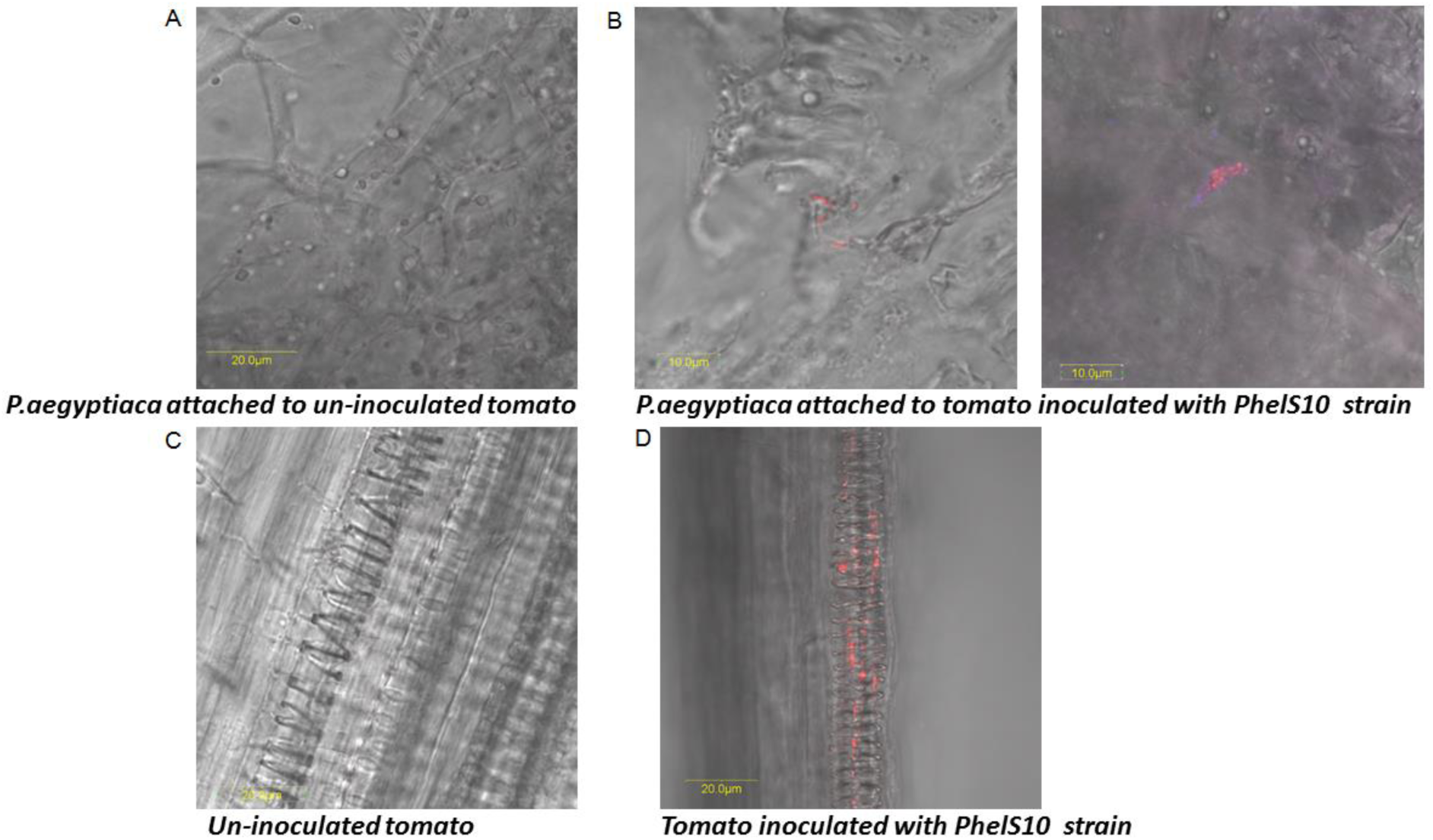
FISH analysis of parasitized and un-parasitized tomato tissue inoculated with *PhelS10* strain. No signal was detected in tomato that were not inoculated with *PhelS10* strain (A+C). This strain was detected in the xylem of tomato inoculated with its culture (red signal, D). In addition, *PhelS10* strain was detected only in broomrape that was parasitized to *PhelS10*-inoculated tomato (red signal, B).

In a previous study (Iasur Kruh et al., 2017), an extensive research was performed examining the microbiota inhabiting both tomato (*Solanum lycopersicum*) host plant and its parasitic *P. aegyptiaca*, using mass sequencing analysis of the 16S rRNA gene. The results showed that bacterial communities of the parasitic weed and its host plant are affected by the parasitism process. Moreover, it was suggested that endophytic bacteria can be transmitted from the parasitic weed to its host and vice versa. The results of the current study (Fig. 3D) confirm the ability of bacteria to move from tomato to its parasitic weed.

Bacterial endophytes, originating from different crops, can be harnessed to control broomrape by inhibiting its growth using the direct mechanism of secreting different secondary metabolites (Joel and Gressel, 2013). If these bacteria can move from their host plant, through the haustorium, and secret their metabolites directly to the parasitic tubercle, this inhibition can be more effective. We suggest that all of the aforementioned traits of the *PhelS10* strain make it a potential efficient biocontrol agent against broomrape in tomatoes.

